# Inhibition of Cardiac p38 Highlights the Role of the Phosphoproteome in Heart Failure Progression

**DOI:** 10.1101/2024.11.20.624554

**Authors:** Sogol Sedighi, Ting Liu, Robert O’Meally, Robert N. Cole, Brian O’Rourke, D. Brian Foster

## Abstract

Heart failure (HF) is a complex condition characterized by the inability of the heart to pump sufficient oxygen to the organs to meet their metabolic needs. Among the altered signal transduction pathways associated with HF pathogenesis, the p38 mitogen-activated protein kinase (p38 MAPK) pathway—activated in response to stress— has attracted considerable attention for its potential role in HF progression and cardiac hypertrophy. However, the exact mechanisms by which p38 MAPK influences HF remain unclear. Addressing knowledge gaps may provide insight on why p38 inhibition has yielded inconsistent outcomes in clinical trials. Here we investigate the effects of p38 MAPK inhibition via SB203580 on cardiac remodeling in a guinea pig model of HF and sudden cardiac death. Using a well-established HF model with ascending aortic constriction and daily isoproterenol (ACi) administration, we assessed proteomic changes across three groups: sham-operated controls, untreated ACi, and ACi treated with SB203580 (ACiSB). Cardiac function was evaluated by M-mode echocardiography, while proteome and phosphoproteome profiles were analyzed using multiplexed tandem mass tag labeling and LC-MS/MS. Our findings demonstrate that chronic SB203580 treatment offers protection against progressive decline in cardiac function in HF. The proteomic data indicate that SB203580-treatment exerts broad protection of the cardiac phosphoproteome, beyond inhibiting maladaptive p38-dependent phosphorylation, extending to PKA and AMPK networks among others, ultimately protecting the phosphorylation status of critical myofibrillar and Ca^2+^-handling proteins. Though SB203580 had a more restricted impact on widespread protein changes in HF, its biosignature was consistent with preserved mitochondrial energetics as well as reduced oxidative and inflammatory stress.

## Introduction

Heart failure (HF) is a complex condition that brings substantial risks to health and life. With over 64 million individuals worldwide affected by heart failure, it has emerged as a pressing concern and reducing its impact has thus become a principal goal(1). The pathophysiology of HF is characterized by a multifaceted interplay of cellular mechanisms, including altered cyclic AMP, cyclic GMP, and Ca^2+^/calmodulin-dependent signaling, Ca^2+^ handling impairment, and mitochondrial oxidative stress. Stress-activated kinases of the mitogen-activated protein kinase (MAPK) family have also garnered significant attention(2, 3). Triggered by osmotic, mechanical, or oxidative stress, MAPK signaling cascades have been implicated as regulators of both cardiac hypertrophy and HF progression(4, 5). MAPKs are a group of highly conserved serine/threonine protein kinases that transmit signals through a multi-level kinase cascade. Four primary subgroups of the MAPK signaling pathway have been recognized: ERK, c-JNK, p38/MAPK, and ERK5(6, 7). These kinases regulate key physiological and pathological processes, including apoptosis and inflammation, as well as proliferation, growth, and differentiation of cardiac resident cells such as cardiomyocytes, fibroblasts, endothelial cells, and macrophages(8).

There are four P38 MAPKs (α, β, ψ, and δ) encoded by genes MAPK14, MAPK11, MAPK12, and MAPK13 respectively. They are activated in response to cellular stressors including oxidative stress, DNA damage, cytokine receptor stimulation(4). Activation leads to phosphorylation of downstream targets including MAPK APK2/3, MSK1 HSP27, and several important cardiac transcription factors (e.g., ATF2, Myc, Stat1, Mef2, Nfat, Creb1, PGC1a)(4, 9). p38 MAPK has been shown to contribute to the growth response of cultured cardiomyocytes to hypertrophic agonists(10),

Of the four p38s, p38α and β have received the greatest scrutiny for their role in HF pathogenesis. Evidence garnered from mouse knockout and over-expression models would indicate that maladaptive p38α activation via phosphorylation by dual-specificity kinases, MKK3 or MKK6, contributes to cardiomyocyte cell death and contractile dysfunction in settings of both chronic pressure overload (11, 12) and ischemia (13), though it may also play an adaptive role in response to acute changes in afterload. P38β expression in the heart is comparatively low (11), although it may have distinct functional roles, for example, in the estrogen-dependent modulation of mitochondrial reactive oxygen species (14).

Targeted P38 α/β inhibition would appear to have therapeutic potential. Both enzymes share high sequence homology, conserved key functional residues within their kinase domains, and have similar pharmacological inhibition profiles. One such inhibitor, SB203580 ([4-(4-fluorophenyl)-2-(4-methylsulfinylphenyl)-5-(4-pyridyl)-imidazole]) exhibits IC50s ranging from 50 and 500 nM, depending on the cell types, which may differ in the relative amount of α and β forms(15–20). Inhibition of p38 MAPK has been proposed as a treatment to inhibit HF pathogenesis(2, 5)

Clinical evaluation of p38 inhibition for several conditions (recovery from myocardial infarction (MI), COPD, or depression) has been pursued, thus far with limited success. Losmapimod, a novel inhibitor of p38 MAPK, was well tolerated upon oral administration and demonstrated efficacy in improving the prognosis of myocardial infarction (MI) patients in a phase II clinical trial (21). However, a larger phase III trial (LATITUDE) showed that, while losmapimod effectively reduced the inflammatory response post-MI compared to placebo, it did not mitigate the risk of major ischemic cardiovascular events (22). Similarly, despite early positive results, a different p38 MAPK inhibitor was recently terminated because it was not expected to meet the primary endpoint in a global Phase 3 trial (REALM-DCM) in patients with symptomatic dilated cardiomyopathy (DCM) due to a mutation of the gene encoding the lamin A/C protein (LMNA)(23). These findings indicate that although p38 MAPK inhibition holds promise in improving cardiac function in the context of heart failure, further investigation of p38 MAPK’s cellular targets and effects is warranted to ascertain the best strategy to optimize interventions that could improve cardiovascular outcomes.

Here, we examine impact of p38 MAPK inhibition with SB203580 (SB) on cardiac HF remodeling in a guinea pig model of HF and sudden cardiac death (24). Our objective is to understand how inhibiting p38 MAPK impacts the progression of HF, focusing on proteome remodeling and alterations in protein phosphorylation associated with HF. We find that p38 MAPK inhibition protects against cardiac decompensation by impacting select classes of HF-associated protein changes while exerting broader protection of the cardiac phosphoproteome.

## Methods

Detailed methods are provided in supplementary file S1-Methods. Methodological references are cited here in the interests of proper attribution (25–36).

## Results

### P38 MAPK inhibition attenuates heart failure progression in guinea pigs

We employed a guinea pig model of heart failure that combines ascending aortic constriction with administration of the β-adrenergic agonist isoproterenol day (1mg/kg/day) via an implanted programmable pump. This model has been validated previously (3, 24). P38 MAPK as demonstrated in **Fig. 1-A,B** is activated by phosphorylation at Thr180/Tyr182 in guinea pig HF, and treatment with SB prevented its activation. The following treatment groups were studied: 1. Sham-operated, serving as Controls; 2. ACi (Aortic Constriction + isoproterenol); 3. ACi-SB (ACi + SB treatment via implanted osmotic pump; 0.5mg/kg/day).

**Figure 1.**
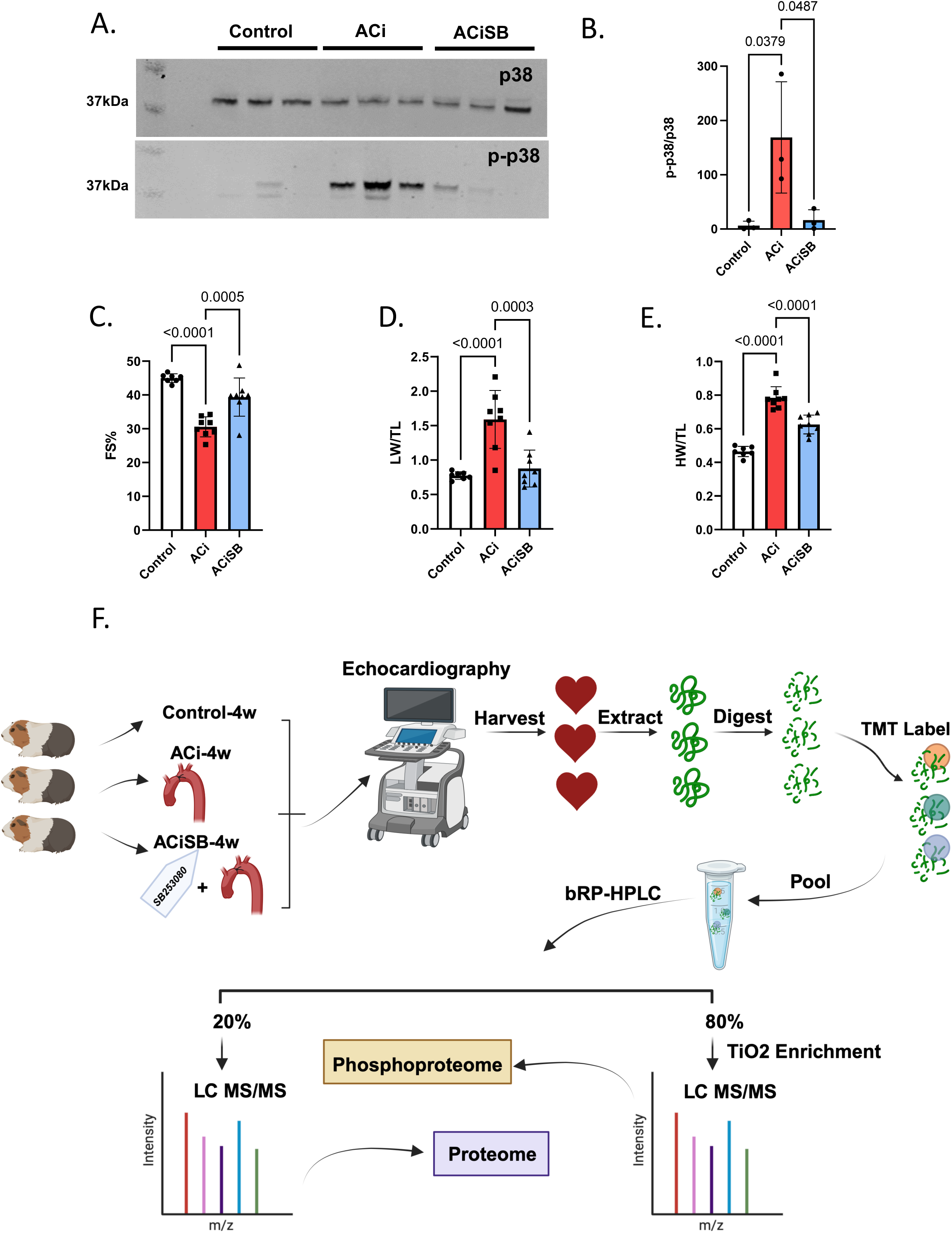
p38 inhibition attenuates heart failure progression in guinea pigs. A. The p38 is activated by phosphorylation at Thr180/Tyr182 in guinea pig heart failure. Treatment with SB203580 prevents activation. B. Ensemble analysis of p38 activation normalized for p38 expression. SB attenuates the decline in Ejection Fraction(C),fractional shortening (D), prevents cardiac hypertrophy (E) and pulmonary edema (F). G. Experimental design-created with BioRender.com.

An appreciable decline in FS was noted in the ACi group compared to its respective Control group (ACi: 30.5 ± 2.9%, n=8; Control: 44.9 ± 1.2%, n=7, p<0.0001), further confirming the validity of our heart failure model. SB treatment effectively abrogated 61% of the decline in FS seen in the ACi-4w group, (i.e. 39.3±5.6%, n=8 vs. 30.5 ± 2.9%, n=8; p-value=0.0005) (**Fig. 1-C**). ACi-induced hypertrophy was also blunted by SB-treatment. Specifically, weight/tibia length was decreased from 0.7±0.06 g/mm (n=8) in the ACi group to 0.6±0.05 g/mm (n=8) in the ACiSB group (p-value<0.001) (**Fig. 1-D**). Additionally, lung weight/tibia length decreased significantly, from 1.5±0.4 g/mm (n=8) to 0.8±0.2 g/mm (n=8) with SB treatment, indicating a reduction in pulmonary edema (p-value=0.0003; **Fig. 1-E**).

To elucidate the impact of P38 MAPK inhibition on the HF proteome, we conducted a comprehensive 16-plex Tandem Mass Tags(TMT) analysis across the three experimental groups. The experimental design is summarized in **Fig. 1-F** for Control, ACi, and ACiSB treated guinea pigs. These samples were extracted, digested, and analyzed as detailed in online supplement S1-Methods. Briefly we used a 2D-LC-MS/MS strategy. TMT-labeled peptides were then pooled prior to high-pH reversed-phase liquid chromatography (bRP-HPLC). The bulk of each concatenated bRP-HPLC fraction (80%) was subjected to titanium dioxide (TiO2) phosphopeptide enrichment. Both enriched and unenriched fractions were subjected to RP-LC-MS/MS. Data analysis consisted of median-sweep scaling, followed by statistical analysis using LIMMA, as we have reported previously (3, 26, 37). Further details regarding chromatography and mass spectrometry apparatus and methods, as well as Ingenuity pathway and STRING functional association network analyses are provided in online methods supplement S1.

### P38 MAPK inhibition partially mitigates protein changes associated with HF

Consistent with our prior work, the ACi protocol elicits substantial proteome remodeling after 4 weeks. Fully 2,480 of 5,016 quantified proteins (i.e. 49%) were differentially expressed in the ACi group (p<0.05 ACi vs Control by LIMMA with post-hoc pairwise contrast). We defined expression as SB-responsive if protein abundance differed significantly between the ACiSB and ACi groups (p<0.05). We found 292 proteins differed between the groups, irrespective of whether they changed significantly between ACi and Control. Thus 227 (of 2,480) proteins whose expression differed from control in the ACi group (i.e. 9%) were deemed SB-responsive. These results are summarized in the Venn diagram of **Fig. 2A**. Complete tabulated protein levels and their statistical analyses are provided in **Supplemental File S2 - Table. Fig. 2B** depicts a PCA biplot of the statistically SB-responsive subset proteins, showing that, even within that subset, the variance of the ACiSB group was distinct from Control, lying between Control and ACi groups. This trend is illustrated more explicitly in the heatmap depicted in **Fig. 2C**, where the protein levels of the SB-responsive group lie between those of the Control and ACi groups. Thus, while select proteins were more SB-responsive than others, on aggregate, SB treatment only partially offset ACi-induced differential protein expression. The volcano plot in **Fig. 2D** highlights some of the proteins whose abundance was most impacted by **SB treatment**.

**Figure 2.**
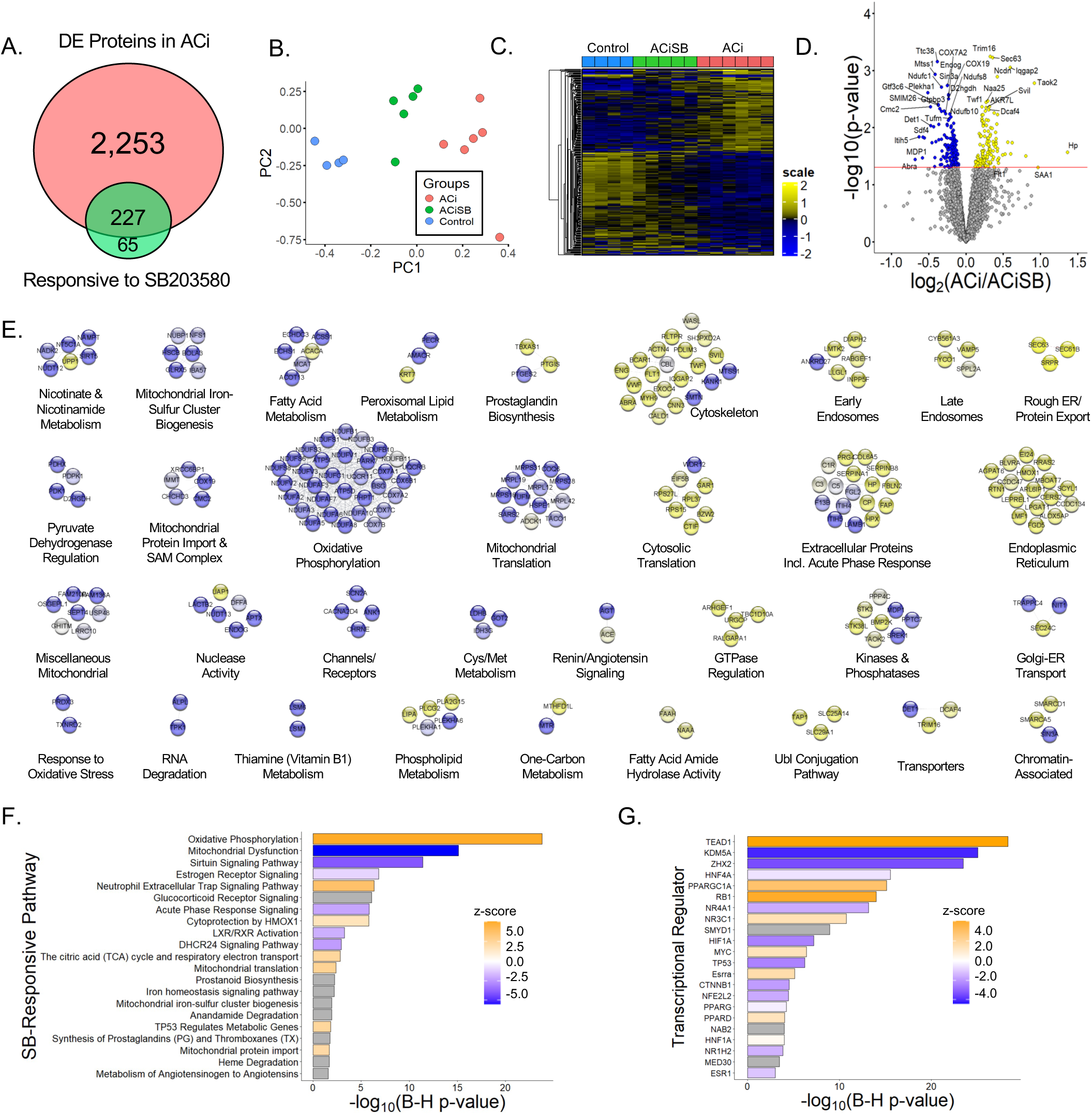
p38 inhibition partially mitigates protein changes associated with HF. A. Venn diagram illustrates that fully 2,480 of the 5,016 quantified proteins change abundance in HF, but only about 1/10 of these proteins are impacted by SB treatment. SB also modulated expression of 65 proteins that were not otherwise changing in HF. B. PCA biplot of proteins changing significantly between ACi and ACiSB. C. Hierarchical clustering of proteins that differed between ACi and ACiSB. D. Volcano plot highlighting specific examples of proteins responsive to SB treatment. E. Functional association network of proteins in D Blue nodes indicate proteins that were downregulated in ACi relative to ACiSB, while yellow nodes indicate protein that were upregulated in relative to ACiSB. F. Pathway analysis of proteins from D. Highlighted pathways were among those with Benjamin-Hochberg-corrected p-values of <0.05. Ochre indicates pathways inferred to be augmented by SB will blue indicates pathways inferred to be attenuated. G. Inferred activation or inhibition of transcription factor activity based on observed protein changes. Color scheme is the same as for F.

For further insights, we subjected 292 proteins to both network- and pathway-based annotation enrichment analyses. The significantly changing proteins in **Fig. 2D** were queried using the STRINGdb functional annotation network. Modules of interest were revealed by Markov clustering in **Fig. 2E**. As in **Fig. 2D**, blue nodes represent proteins downregulated in ACi that were SB-responsive while yellow nodes indicate responsive upregulated proteins. The 33 modules depicted encompass 214 of the 292 SB-responsive proteins and summarize the major features of the dataset. Among downregulated, yet SB-responsive proteins, several modules correspond to mitochondrial processes or pathways, including oxidative phosphorylation, mitochondrial translation, mitochondrial iron-sulfer cluster biogenesis, mitochondrial protein import and nicotinamide metabolism. Proteins of the respiratory complexes were particularly SB-responsive (also see **Fig 4**). Among the upregulated proteins, major nodes implicate extracellular and acute phase response proteins, ER and endosomal proteins, and proteins of the cytoskeleton. Acute phase response proteins were among the most SB-responsive (also see **Fig 5**).

Complementary Ingenuity pathway analysis **(Fig. 2F)** is consistent with network-based annotation, implicating both oxidative phosphorylation and acute phase response signaling, whose dysregulation in the ACi group was ameliorated by treatment with SB. Finally, Ingenuity upstream regulator analysis (URA) provides a set of candidate transcription factors whose activity might explain the changes in observed protein levels arising from SB treatment **(Fig. 2G).** URA strongly implicates Tead1, whose activity could explain the coordinate expression of 20 respiratory complex proteins, particularly from complex I (see **Fig. S2A**). Prior work has shown that Tead1 deletion decreases phospholamban phosphorylation, SERCA2a expression, and mitochondrial gene expression, resulting in cardiomyopathy in mice (38). Here, the inferred involvement of Tead1 is consistent with reports that stress-induced activation of p38 MAPK results in its interaction with, and phosphorylation of, Tead1 in the cytoplasm, inhibiting its nuclear function as a transcription factor in the Yap/Taz pathway(39). The Tead1 transcriptional program partially overlaps with Rb1 and PGC1α programs. Together Tead1, Rb1 and PGC1α regulation likely account for most of the mitochondrial protein down regulation in ACi that responds to SB (**Fig. S2B**).

With respect to mechanisms of chromatin remodeling, Kdm5a, a lysine-specific demethylase involved in the regulation of gene expression is inferred to be strongly inhibited by SB-treatment (p = 1.3×10^−11^, z-score= −4.1). Kdm5a has previously been identified as a key regulator of cardiac fibrosis and is upregulated in fibroblasts from patients with dilated cardiomyopathy (DCM) via the angiotensin II and PI3K/AKT signaling pathways(40).

### Expression of MAPK cascade proteins change in HF, but are largely unresponsive to SB

MAPK cascade proteins including Mapk14 (p38a), Mapk9 (Jnk2), Mapk1 (Erk2), along with their upstream activators, Map2k1, Map2k2, Map2k3 and Map4k5 (Khs1), exhibited alterations in their abundances in ACi but showed no significant responsiveness to SB treatment **(Fig. 3).** An exception to the trend, Map3k17 (Taok2), which is an upstream kinase in the p38 MAPK cascade, didn’t change in ACi but was decreased with SB treatment. Notably, while the abundance of Mapk14 (p38α) and Map4k5 (Khs1) did not undergo significant changes, their phosphorylation was markedly altered, as depicted in **Fig. 7** and discussed hereinafter. With respect to known p38 MAPK substrates, the expression of most remained unchanged in ACi. Notably, among the substrates of p38, only Gys1, Mef2d, and Spag9 and Lsp1 exhibit altered abundances in ACi, furthermore only Lsp1 was responsive to SB treatment (**Fig. S3**).

**Figure 3.**
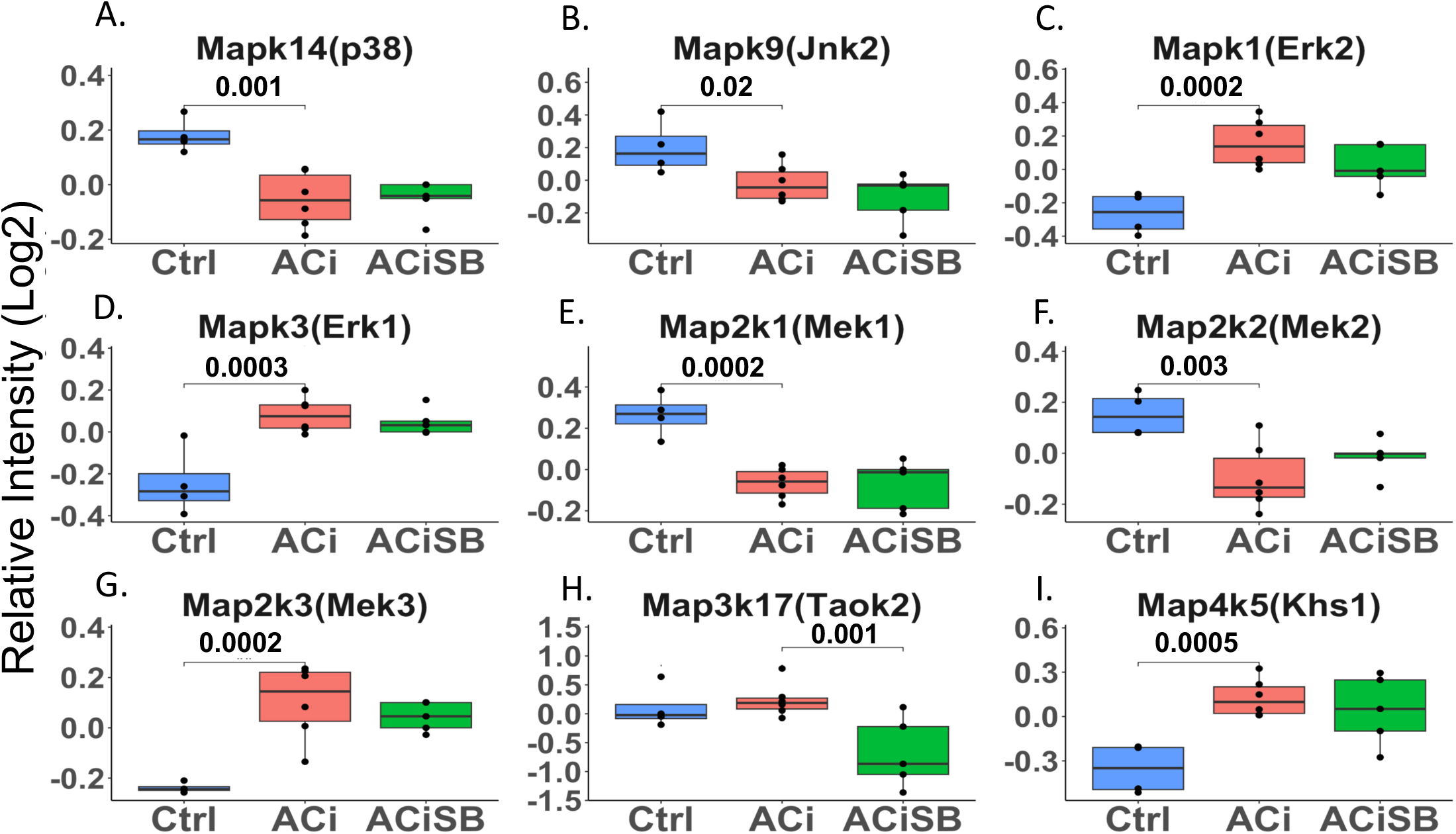
Several MAPK Cascade proteins are differentially expressed in heart failure, but largely unresponsive to SB. Global significance (p < 0.05) established via F-test, with p-values derived from LIMMA contrast matrices for inter-group comparisons. P-values < 0.05 are indicated by faceted brackets.

### SB203580 Curbs Changes in Mitochondrial and Acute Phase Response Proteins

Downregulation of mitochondrial proteins is a consistent biosignature of HF and correlates with mitochondrial dysfunction. The mitochondrial network modules highlighted **in Fig 2E** are consistent with prior studies. **Fig 4** specifically shows that, in particular, several subunits of respiratory complexes I (Ndufa8, Ndufaf7, Ndufb1, Ndufc1, Ndufs1, Ndufv1) and IV (Cox6b1, Cox7C, Cox11, Cox19) were downregulated in the ACi group. **Fig. 4** also shows that their decline is substantively abrogated in the ACiSB group. Atp5f1d, a subunit of ATP synthase, and Uqcrb, from complex III, are likewise SB responsive. The Pdks, or pyruvate dehydrogenase kinases, are key regulators of pyruvate metabolism to acetyl-CoA via phosphorylation of the pyruvate dehydrogenase complex. In HF, Pdk1 levels typically decline while Pdk4 levels rise. This observation holds in the ACi model. SB-treatment had a mild, though significant, impact on Pdk1, although there was no significant effect on Pdk4. Taken together, this suggests that SB treatment might offset impaired mitochondrial function in HF.

**Figure. 4.**
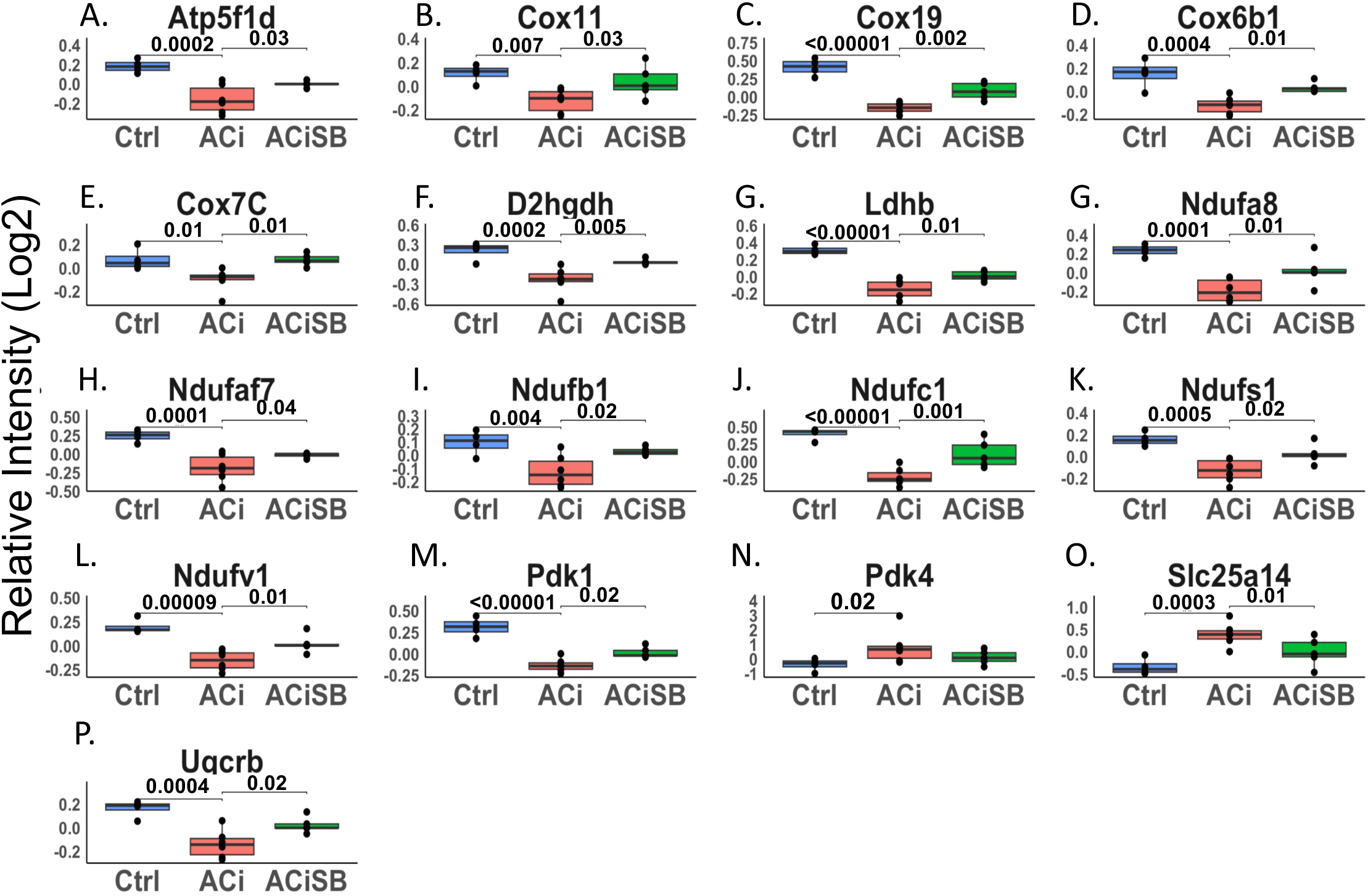
Expression of many proteins involved in mitochondrial bioenergetics are changing significantly and are responsive to SB. Global significance (p < 0.05) established via F-test, with p-values derived from LIMMA contrast matrices for inter-group comparisons. P-values < 0.05 are indicated by faceted brackets.

Like mitochondrial dysfunction, inflammation and activation of the acute phase response is a hallmark of human HF and recapitulated here in the guinea pig ACi Model. Several proteins associated with innate immunity were significantly up regulated in ACi (**Fig. 5**), and all these trended towards mitigation of the response in the ACiSB group. However, owing to high variation in the response for this class of proteins, only 4 of these showed statistically significant inhibition by SB treatment; specifically, Hp, Serpina1, lqgap2, and Vwf **(Fig. 5)**.

**Figure. 5.**
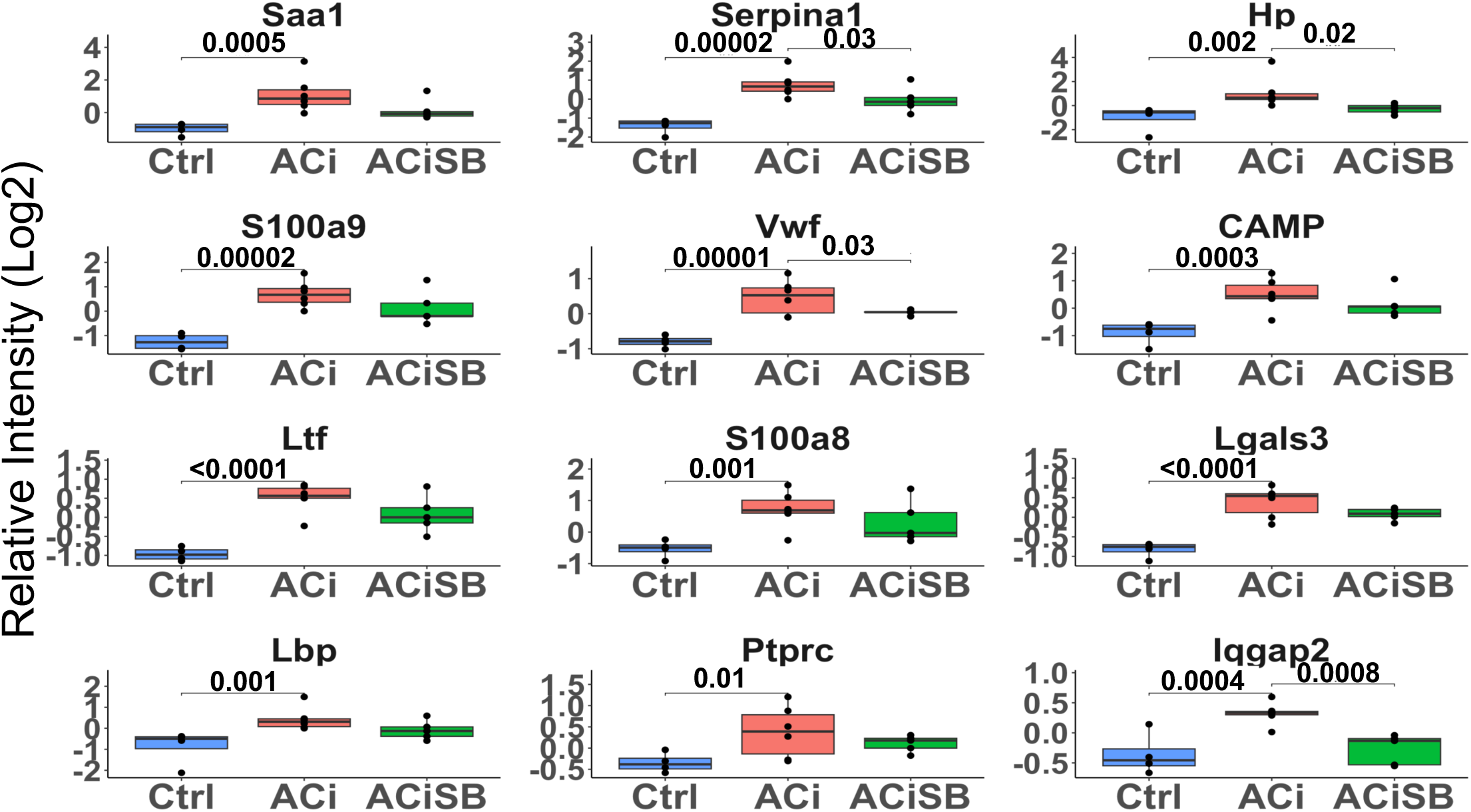
Expression of many acute phase proteins are significantly increased with ACi. Out of those that are changing Hp, Serpina1, Iqgap2 and Vwf are SB-responsive. Global significance (p < 0.05) established via F-test, with p-values derived from LIMMA contrast matrices for inter-group comparisons. P-values < 0.05 are indicated by faceted brackets.

### Characteristics of the Cardiac Phosphoproteome

Our study identified 4,310 unique high confidence phosphopeptides (1% FDR), encompassing 3,844 unique phosphorylation sites. 3,482 phosphopeptides (80%) could be linked to a quantified protein. Of the 5,016 proteins quantified, phosphorylation sites were detected for 1,129 (22%) of them. Phosphoproteome analysis of the failing heart is complicated by the fact that nearly half of all quantified proteins in our expression proteome are differentially expressed between ACi and Control groups. As **Fig. 6A** illustrates, the observed change in phosphorylation levels strongly correlate with changes in the levels of underlying protein between the Control and ACi groups (Pearson r = 0.72). More explicitly, the R^2^ value of 0.52 indicates that fully half of the variance in measured phosphorylation can be attributed to changes in underlying protein abundance. To discriminate between changes in *bona fide* phosphosite occupancy from changes arising from differential phosphoprotein abundance, phosphopeptide signals were normalized to protein levels, where possible. Normalization was performed by subtracting logged & median-swept relative protein abundances from logged & swept phosphopeptide abundances, for each biological replicate. Following normalization, changes in phosphopeptide abundance showed only a mild residual inverse dependence on changes in protein abundance (R^2^ = 0.08; **Fig. S4**).

**Figure 6.**
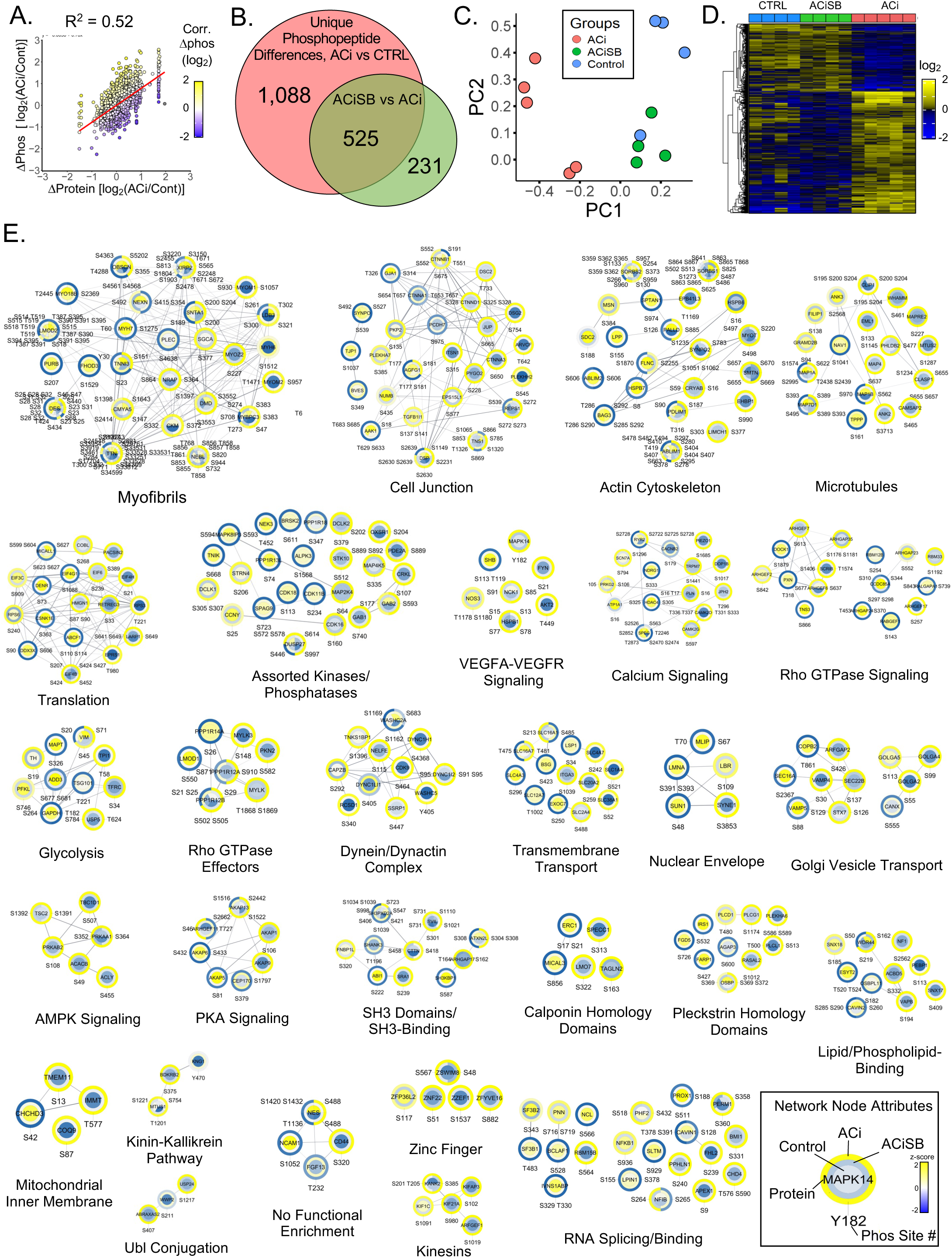
p38 inhibition attenuates substantial changes in phosphorylation associated with HF. A. Scatterplot reveals the correlation between changes in phosphosite changes in HF relative to changes in underlying protein abundance. Just over 50% of the variance in phosphorylation can be explained by changes in relative protein levels. B. Venn analysis reveals that of the 1613 unique phosphosites changing between control and ACi, 32% of were impacted by SB treatment. A further 231 phosphosites differed between ACiSB and ACi, despite no significant change in ACi relative to controls. C. PCA biplot analysis of the 525 unique phosphopsites, indicate the character of the SB-significant phosphoproteome is closer to that of control hearts than failing hearts. D. Hierarchical cluster highlights that about the largest impact of SB on the phosphoproteome was by inhibiting phosphorylation that otherwise increases in failing hearts. E. Depicts the z-scored phosphosite signals, superimposed on a functional annotation network. Functional modules are revealed through network Markov clustering using the String score for edge-weighting. A legend of network node attributes is provided.

### p38 MAPK inhibition impacts a major portion of HF-associated phosphorylation changes

Statistical analysis was conducted on all phosphopeptides. 1,613 changed significantly between Control and ACi groups, of which 525 (33%) were significantly impacted by SB treatment. A further 231 phosphopeptides differed between ACiSB and ACi, irrespective of whether they changed in ACi relative to Controls (**Fig. 6B**), PCA biplot analysis of the SB-responsive phosphosites is illustrated in **Fig. 6C** (p<0.05, ACi vs ACiSB). This is illustrated explicitly in the hierarchically-clustered heatmap in **Fig. 6D**, where three major SB-dependent trends can be distinguished. First, there is a set of sites that become hyperphosphorylated in ACi that are largely prevented by SB treatment. Secondly, there are phosphorylated sites that become hypophosphorylated in ACi but which SB-preserves at Control levels, perhaps through indirect impact on intermediary phosphatases. Thirdly, SB inhibits phosphorylation of a unique set of phosphorylation sites that are phosphorylated in both Control and ACi conditions.

To extract greater insight into processes impacted by SB, the quantitative information from the heatmap in **Fig. 6D** was superimposed onto a Markov-clustered STRINGdb functional annotation network to visualize changes in relative phosphosite occupancy in context of ontologically-enriched modules (**Fig. 6E).** As depicted in the legend **(6E, bottom right)** The center of the node denotes Control levels of phosphorylation, the outer ring represents phosphorylation levels in ACi and the middle ring denotes phosphorylation in the ACiSB group.

Among phosphoproteins, cytostructural proteins constitute a major proportion. Large modules include the myofibrils, cell junction, actin cytoskeleton and microtubules. Additional modules consist of cytoskeletal regulatory proteins such as the Rho GTPases and Rho GTPase effectors. A second broad category of phosphoproteins encompasses membrane-associated or membrane-trafficking ontologies including Golgi vesicle transport, dynein/dynactin complexes, kinesins and nuclear envelope proteins. Channels and transporters are encompassed by the calcium signaling and transmembrane transport modules. Finally, a third major category represented in **Fig 6E** are proteins involved phosphorylation-mediated signal transduction. Notable modules include Protein kinase A (PKA) signaling, AMP Kinase (AMPK) signaling, Vascular endothelial growth factor (VEGF) signaling, assorted kinases and phosphatases, as well as SH3 domains/binding.

### Phosphorylation of p38α, upstream kinases, and downstream substrates are responsive to SB

MAPK family signaling is characterized by extensive cross-talk and feedback regulation by both direct phosphorylation and through indirect phosphorylation/dephosphorylation through intermediary kinases and phosphatases, as well as indirect effects on the gene regulation of intermediary phosphatases. Accordingly, one might expect SB to affect the phospho-status of upstream regulators of p38 Mapk, Erks and Jnks, as well as their known substrates p38 Mapk substrates. **Fig 7** Indicates that Mapk14 (p38α) showed a significant decrease in phosphorylation upon SB treatment, as shown earlier by western blot (**Fig 1A**). Map2k4(Mek4), Map3k7(Tak1), Map4k5(Khs1), and Map4k6(Mink1) displayed increased phosphorylation in the ACi group, which, for Map2k4, was significantly attenuated by SB treatment. Conversely, Map3k2 (Mekk2) showed a significant decrease in phosphorylation in the ACi group.

**Figure 7.**
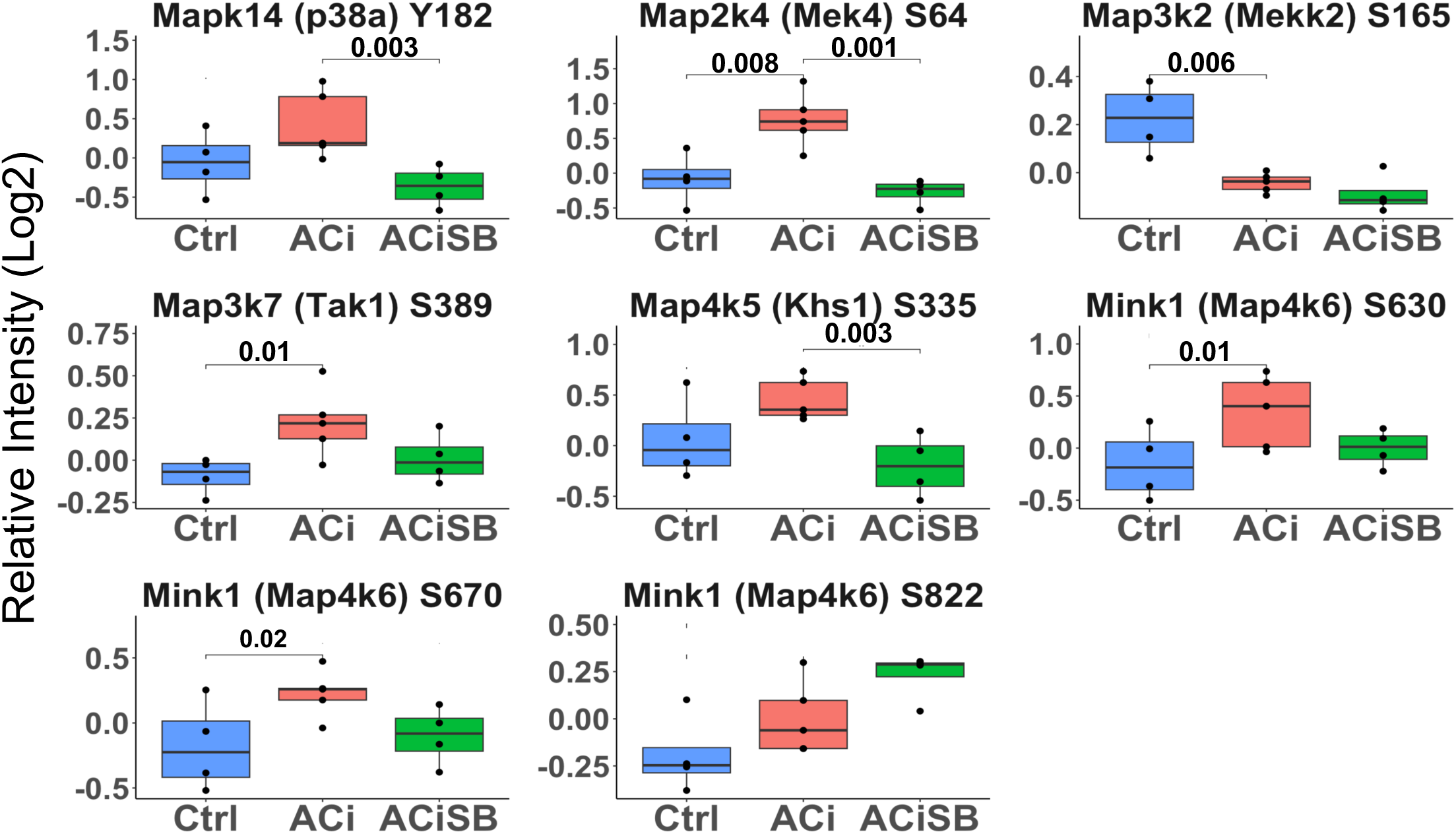
Phosphorylation of p38a and Several Upstream Kinases are Responsive to SB. Global significance (p < 0.05) established via F-test, with p-values derived from LIMMA contrast matrices for inter-group comparisons. *p*-values <0.05 are marked with brackets.

With respect to known protein substrates of p38 Mapk, phosphorylation levels of Spag9 (Ser 584) and Nelfe (Ser S115) increased with HF, and exhibited significant responsiveness to SB treatment. Additionally, Hspb6 (Ser 16; Hsp20) stood out as SB sensitive. This site increased by ∼2.5-fold in the ACi group compared to Controls, which was mitigated by SB treatment **(Fig S6)**.

### Phosphorylation of select ion transport proteins sensitive to SB

Phosphorylation status of several ion transport channels, including Atp1a2, Cacna1c, Cacnb2, Kcnh2, Kcnq1, Trpm7, Piezo1, and Clcc1 were also altered in the context HF; Only Trpm7, Prezo1, Clcc1 were responsive to SB treatment. **(Fig 8).**

**Figure 8.**
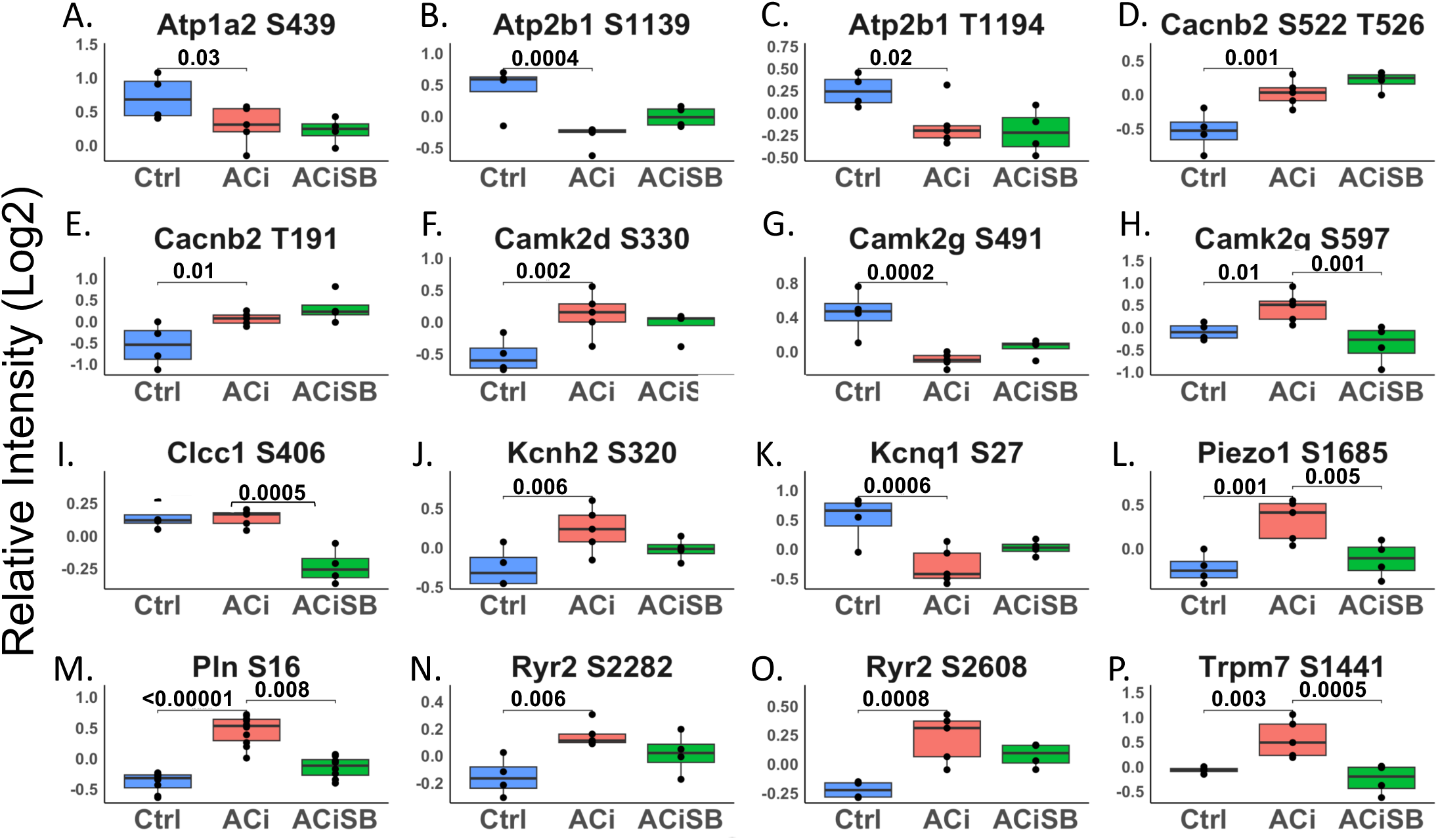
Phosphorylation of select proteins of the ion transport are sensitive to SB. Global significance (p < 0.05) established via F-test, with p-values derived from LIMMA contrast matrices for inter-group comparisons. *p*-values <0.05 are marked with brackets. Only a subset of results is shown in the figure.

## Discussion

In this study, we showed that inhibition of p38 MAPK with SB203580 offered considerable efficacy in protecting against experimental HF, offsetting the bulk of the decline in fractional shortening, as well as alleviating both pulmonary edema and cardiac hypertrophy. Our proteomic studies indicate that this protection is characterized by a substantial impact on the observable phosphoproteome. The scope of the impact on the proteome was modest by comparison, though the impacted processes, most notably mitochondrial function and inflammation, are key determinants of cardiac function. Taken together, notwithstanding the broad protein expression changes in HF, the data are consistent with a contributing role for p38 MAPK-driven phosphoproteome modifications in cardiac decompensation. We discuss the impact of SB203580 on the proteome and the phosphoproteome, in turn.

### p38 MAPK inhibition offsets the impact of ACi on the mitochondrial proteome

Though scarcely 10% of differentially expressed proteins in HF were deemed SB-responsive, many were associated with mitochondrial pathways. The protein-level data suggest that SB-treatment maintains mitochondrial functional integrity by preserving the stoichiometry of the respiratory oxidative phosphorylation (oxphos) complexes. Not only does SB prevent the decline in levels of over 30 complex subunits, it offsets declines in the mitochondrial protein translation machinery that make the mitochondrially encoded subunits and normalizes the expression of proteins involved in iron-sulfur biogenesis and complex assembly. SB further impacts levels of the MICOS complex members responsible for cristae formation, thereby maximizing bioenergetic efficiency. By maintaining respiratory chain integrity, SB may serve to optimize ATP production, while minimizing mitochondrial ROS generation, a principal driver of HF pathogenesis(3). We further note that key antioxidant defense proteins including thioredoxin reductase 2 (TrxnRd2) and ferredoxin reductase were also SB-responsive. Finally, SB may help to preserve proper mitochondrial substrate utilization by ameliorating HF-induced changes in fatty acid oxidation enzymes and perhaps forestalling the switch in pyruvate dehydrogenase kinase activity. Recent research suggests Pdk4 inhibition as a promising strategy for HFrEF therapy. Targeting Pdk4 could offer a novel adjunctive therapy for HF, especially for patients resistant to conventional treatments.(41)

Potential mechanisms for preserving mitochondrial integrity and fatty acid oxidation are suggested by the URA analysis, which implicated the lysine demethylase Kdm5A and the transcription factors Tead1, PGC1a and Rb1. The network diagram in **Fig S2B** shows that these four transcriptional regulators could reasonably account for the coordinate expression of mitochondrial and metabolic proteins. Tead1 plays a vital role as a central transcriptional hub, autonomously regulating a broad network of genes associated with mitochondrial function and biogenesis (42). Ablating *Tead1* in mice causes cardiomyopathy(38). Perhaps its best documented role is as one of the end-effector transcription factors of the HIPPO-Yap/Taz pathway, where it is activated by the nuclear translocation of dephosphorylated Yap/Taz (43). However, p38 has also been shown to downregulate Tead1 activity directly, independently of Yap/Taz. Specifically, p38 phosphorylates Tead1, prevents its shuttling from the cytoplasm to the nucleus (39, 44). Applied here, p38 hyperactivation would then be expected to inhibit the Tead1 bioenergetic program leading to mitochondrial dysfunction. SB-treatment, by inhibiting Tead1 phosphorylation, could ensure there is no brake on nuclear shuttling, and thus preserve Tead1-mediated transcription.

### SB203580 curbs the Acute Phase Protein Response in HF

The P38 MAPK pathway regulates inflammatory cytokine expression, immune cell functions, and cardiac healing(45), Acute inflammation is crucial for cardiac protection, but unresolved inflammation can lead to heart failure(46). In our study, SB treatment reduced acute phase proteins and inflammation in heart failure, likely contributing to improved cardiac function. Because the acute phase proteins identified are typically ones secreted into the circulation by the liver in response to inflammatory cytokines, part of the SB beneficial effect in HF is likely to be linked to inhibition of systemic inflammation.

### SB203580 Broadly Impacts the HF Phosphoproteome

We identified over 800 phosphopeptides that were responsive to SB treatment. Notwithstanding the specificity of the inhibitor, the set of SB-responsive phosphosites extends to peptides that don’t conform to the canonical MAPK family consensus site characterized by a proline residue immediately C-terminally adjacent to the phosphosite (S/T-Pro). Presumably, the 4w administration of SB influenced the phosphostatus of both direct substrates, as well as indirect targets, by influencing the expression and/or activity of other kinases and phosphatases. This was keenly apparent in **Fig 6E**, where several modules within the SB-responsive phospho-network were associated with phosphorylation pathway signaling by other kinases and phosphatases (e.g. VEGF, PKA and AMPK substrates).

The impact of SB treatment on PKA signaling is noteworthy, given that HF is characterized by a loss of β-adrenergic responsiveness, manifested as a reduction in cardiac contractile power and slowed ventricular relaxation. A hallmark of the PKA signaling deficit is the progressive dephosphorylation of the thin filament regulatory protein, cardiac Troponin I (cTnI) at serines 23 and 24 (database numbering). Dephosphorylation aberrantly increases the Ca^2+^-sensitivity of myofibril contraction and slows myofibril relaxation rate. Here, we show that SB treatment indirectly preserves cTnI Ser23 phosphorylation. cTnI is also phosphorylated at Ser150 (Ser151 herein) by AMPK, which has been shown to blunt PKA phosphorylation at Ser 23/24. We also demonstrate that Ser151 phosphorylation is elevated in HF, but is blunted by SB-treatment. Thus, an unexpected or emergent consequence of chronic SB treatment in HF is that it preserves β-adrenergic signaling to the thin filaments. Besides cTnI, SB-treatment impacts phosphorylation of over 25 myofibrillar proteins on nearly 100 unique phosphopeptides, the majority of which have not been characterized. In addition to contractile proteins, the myofibrillar substrates include Z-disk proteins and structural links to the costameres. We therefore speculate that SB-treatment could also modulate mechanotransduction from the sarcolemma to the myofibrils and the nucleus.

How p38 inhibition preserves β-adrenergic signaling homeostasis is unclear, though we observed substantially altered phosphorylation among the A-kinase anchoring proteins or AKAPs, including AKAP1, AKAP5, AKAP6, AKAP9, and AKAP13 that localize PKA and/or PKC signaling to discrete nanodomains. The AKAPs are particularly notable, as several have documented roles in Ca^2+^ handling, serving as docking points for PKA, which in turn phosphorylate and modulate the activity of ion channels. Examples include the role of AKAP5 in regulation of Ca^2+^-influx through the T-tubular Cav1.2 and the role of AKAP6 in regulation of sarcoplasmic reticulum Ca^2+^ release through Ryr2. Coincidentally, we note that SB-normalizes phosphorylation of the Cav1.2 regulatory channel (Cacna1b), as well as two mechanosensitive sarcolemmal divalent cation channels, TrpM7 and Piezo1. SB likewise preserved the phosphorylation state of Ryr2 and phospholamban.

Apart from PKA signaling, we also note that SB treatment impacted AMPK signaling and Ca^2+^-Calmodulin activated kinase (CamKII) phosphorylation. Specifically, SB abrogated the ACi-induced hyperphosphorylation of the AMPKa1 catalytic subunit (Prkaa1) at Ser351 (guinea pig numbering; equivalent to Ser496 in the mouse), within its AMP sensor domain. Phosphorylation of Ser496 by either PKA or Akt suppresses AMPK activity. SB similarly prevents AMPK beta subunit2 (Prkab2) hyperphosphosphorylation at Ser108, within its glycogen-binding domain, which would be predicted to impair glycogen binding. Hyper activation of the Ca^2+^-Calmodulin activated kinases, particularly Camk2D is well documented in HF. The activation process is mediated, in part through autophosphorylation at multiple sites, some of which are better characterized than others. Here we show phosphorylation at Thr337, which has previously been shown to increase kinase activity, is blunted by SB treatment.

### Comparing the impact of SB203580 and the antioxidant mitoTEMPO on HF progression

Our previous research highlighted the significance of targeting mitochondria as a therapeutic approach for heart failure. Employing the mitochondrially-targeted antioxidant MitoTEMPO normalized cellular ROS levels. Additionally, administering MitoTEMPO to HF animals in vivo prevented and reversed HF, mitigated the risk of sudden cardiac death by reducing repolarization dispersion and ventricular arrhythmias, attenuated the chronic HF-induced remodeling of proteome expression, and prevented specific alterations in the phosphoproteome(3). Furthermore, the MAPK kinase pathways emerged as a pathway sensitive to mitochondrial ROS (mROS), known to be activated by it and exhibiting altered signaling in HF models. Notably, the activation of MAPK was evident in the subset of proteins displaying changes in the expression proteome of failing hearts, a phenomenon moderated by MitoTEMPO treatment.(3).

Even though MitoTEMPO’s effect on protein remodeling in heart failure was significantly greater than that of SB and had a wider influence, comparing the effects of MitoTEMPO and SB on the HF proteome highlights the diverse remodeling pathways triggered by mitochondrial ROS, including the activation of p38 MAPK. It is noteworthy that p38, downstream of mitochondrial ROS, plays a role in oxidative phosphorylation and mitochondrial processes, suggesting that its inhibition could yield beneficial effects.

### Limitations of the study

We have captured a snapshot of how SB treatments impact HF phosphoproteome in HF pathogenesis. Therefore, it is a challenge to discern how many of the SB-responsive phosphopeptides are *bona fide* p38 MAPK substrates. While the presence of a proline at P+1 is a defining feature of the p38 MAPK consensus sequence, assigning substrates is complicated by the fact that the consensus sites of other “proline-directed” kinases (Mapks, Cdks) are highly similar(47). Moreover, though a subset of proline-containing SB-responsive sites represent p38 substrates, the experimental design (i.e. chronic SB treatment) makes it difficult to parse the direct p38-mediated impact of SB on substrate phosphorylation. Only a time-course of SB-mediated p38 inhibition in cardiac cell types would help address the issue. Finally, we are using whole heart lysates, which do not distinguish between effects on different cell types, such as fibroblasts, whose activation state is known to be modulate by p38 MAPK (48). This will require additional cell fractionation studies in future work.

## Conclusion

In conclusion, chronic SB treatment elicits substantial protection against HF, by exerting effects on the phosphoproteome that percolate beyond direct inhibition of p38 to influence the broader web of cardiac kinase signaling from PKA to AMPK and CAMKII, ultimately ameliorating the phospho-status of key myofibrillar and Ca^2+^-handling substrates. The impact of SB on the underlying HF proteome, while not expansive, is consistent with preserved energetics as well as reduced oxidative and inflammatory stress. Further research and clinical investigation are warranted to unravel the full potential of targeting p38 MAPK as a part of therapeutic strategies aimed at improving outcomes in HF management.

## Data Accessibility & Reproducibility

The MS proteomics raw data (.raw), complete search results (.msf), and spectra (.mzidentML) have been deposited to the ProteomeXchange Consortium (http://www.proteomexchange.org/;(49)) via the PRIDE partner repository (50) with the data set identifier PXD058012 and 10.6019/PXD058012. The R code used for data analysis along with macros used for finding homologous human peptides can be found at https://github.com/Frostman300/p38-upload

---

**Figure. S1.**
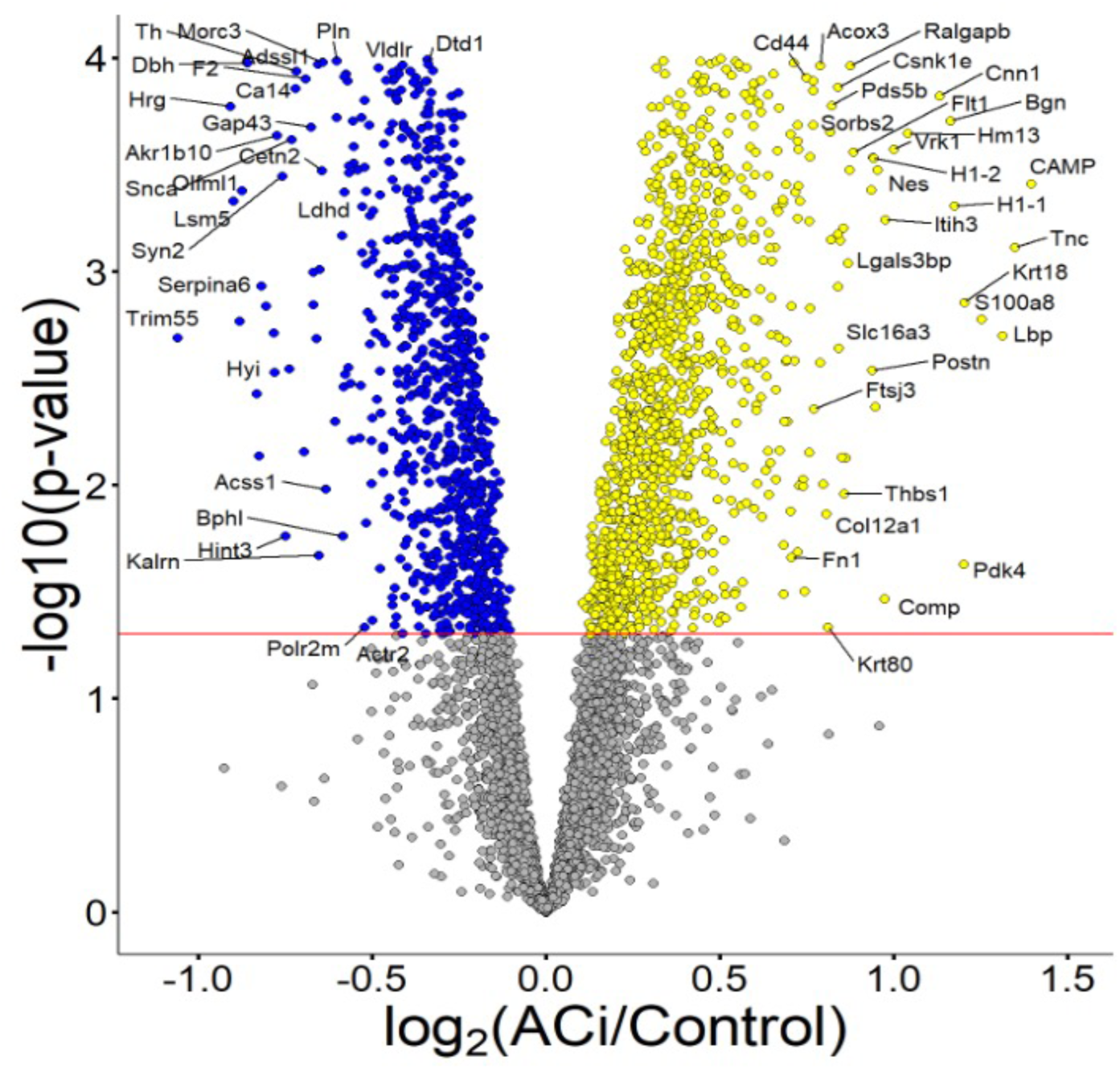
Volcano plot highlighting specific examples of proteins changing between ACi and Control. yellow nodes demonstrate proteins that were significantly upregulated in ACi vs Control; Blue nodes demonstrate proteins that were significantly downregulated in ACi vs control

**Figure. S2.**
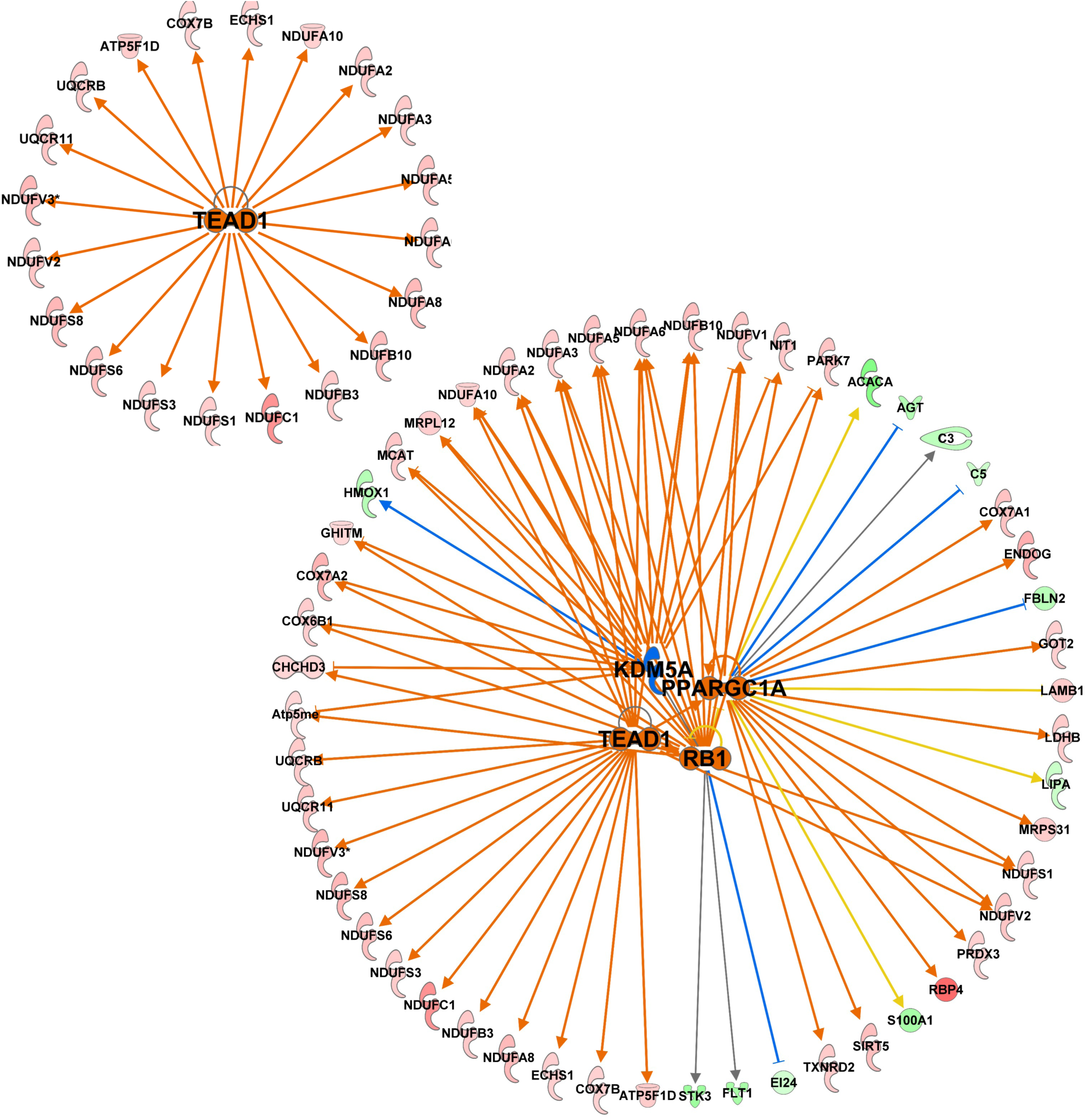
A network diagram of transcriptional regulators, including TEAD1, KDM5A, PPARGC1A, and RB1 previously shown in Figure 2.G.

**Figure S3.**
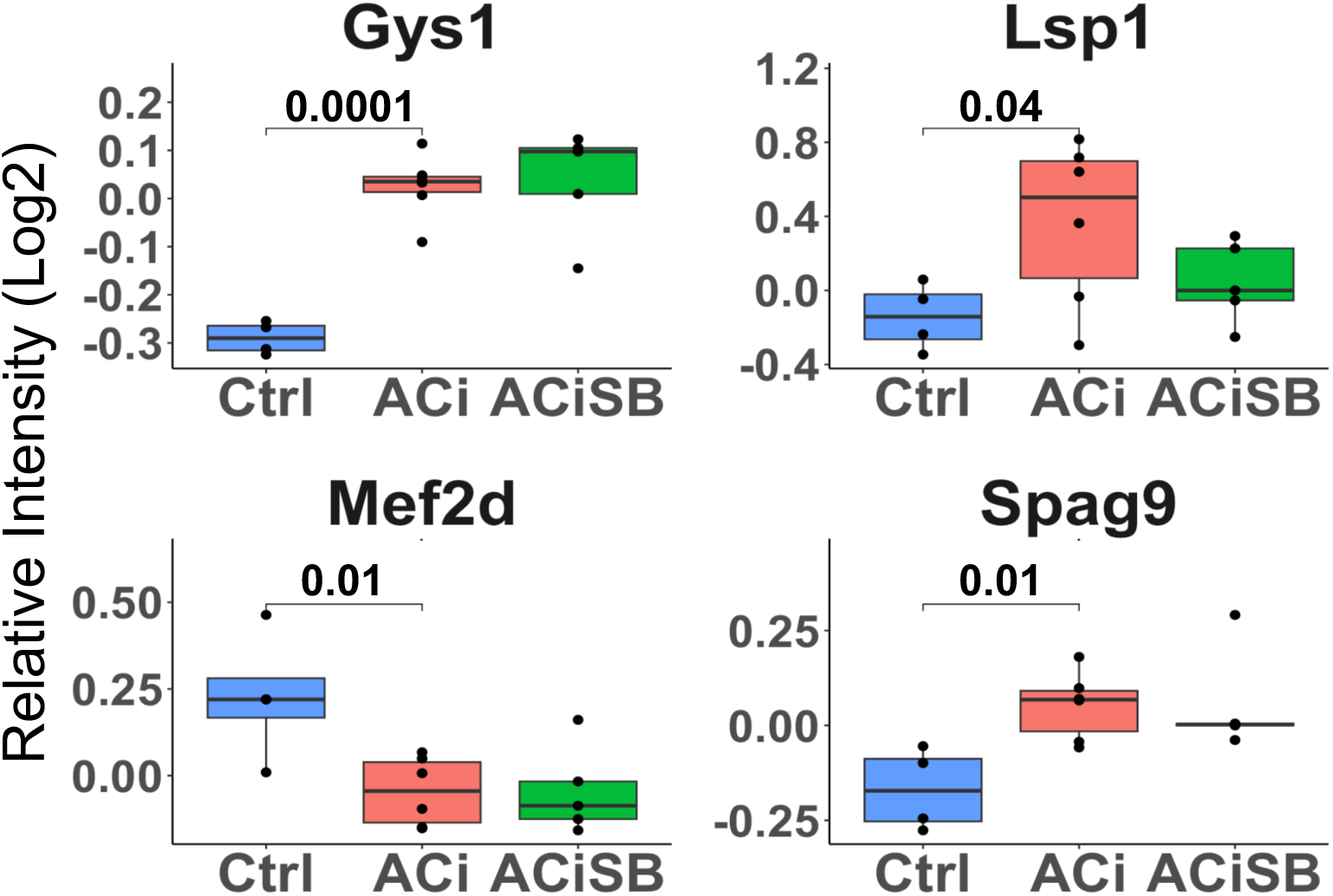
Expression of many p38 MAPK substrates does not change in Aci. Global significance (p < 0.05) established via F-test, with p-values derived from LIMMA contrast matrices for inter-group comparisons. P-values < 0.05 are indicated by faceted brackets.

**Figure S4.**
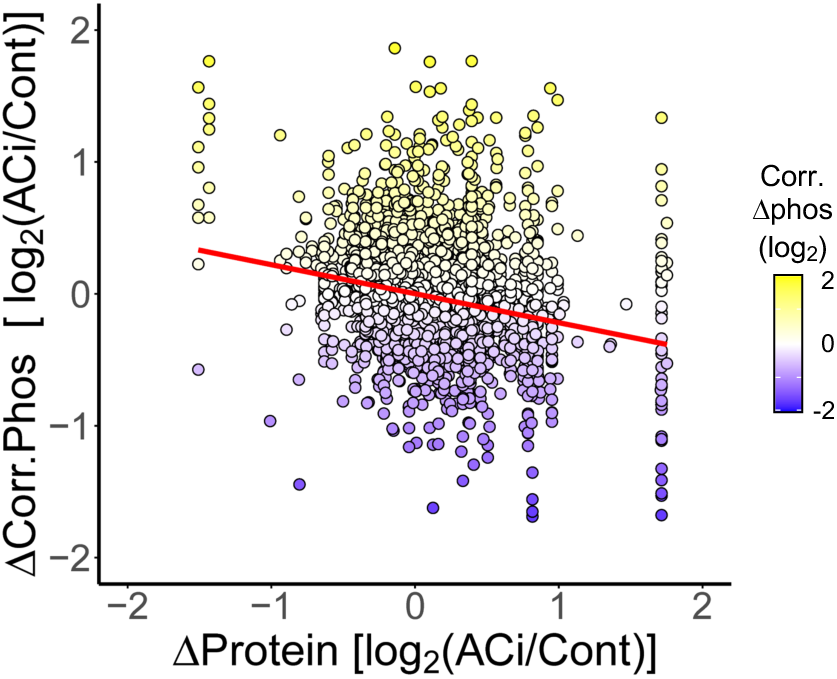
Correcting phosphorylation signals for underlying protein abundance substantially reduces the dependence of phosphorylation on protein dynamics in HF.

**Figure S5.**
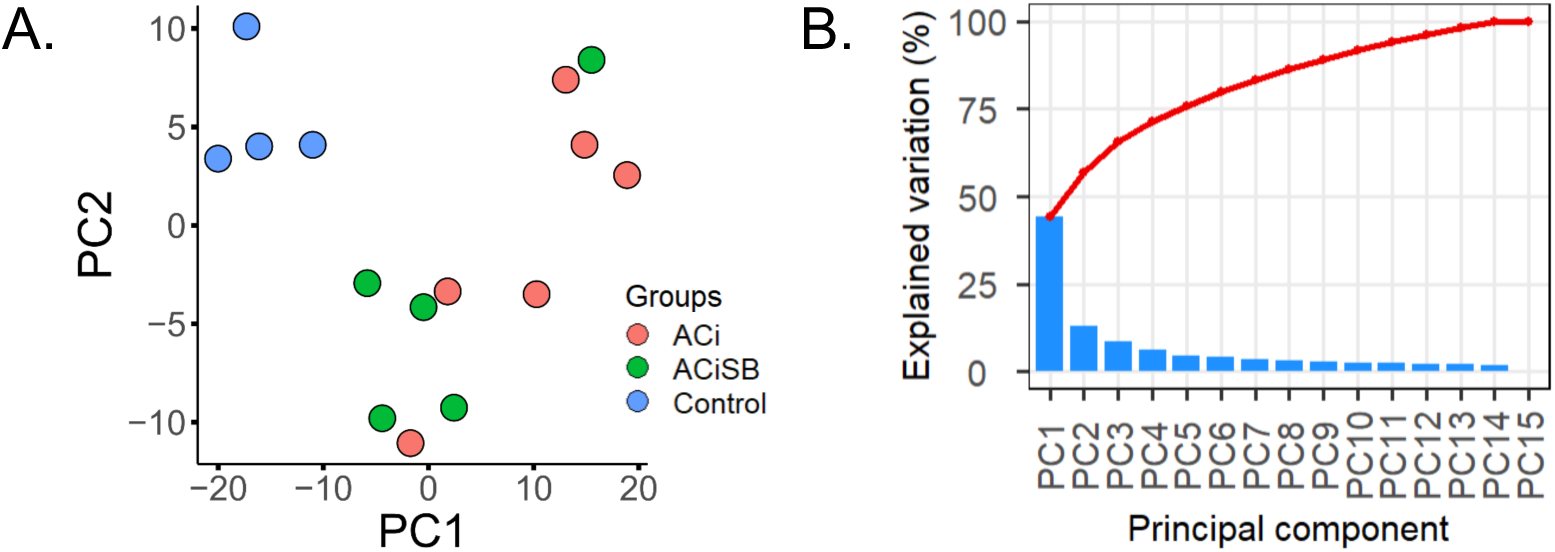
Phosphoproteome, uncorrected for protein abundance. A. PCA biplot indicates of the ACi phosphoproteome from control. SB mitigates remodeling, though changes in protein abundance accounts for a large measure of the variance. B. Scree plot indicating the contribution of each principal component to experimental variance.

**Figure S6.**
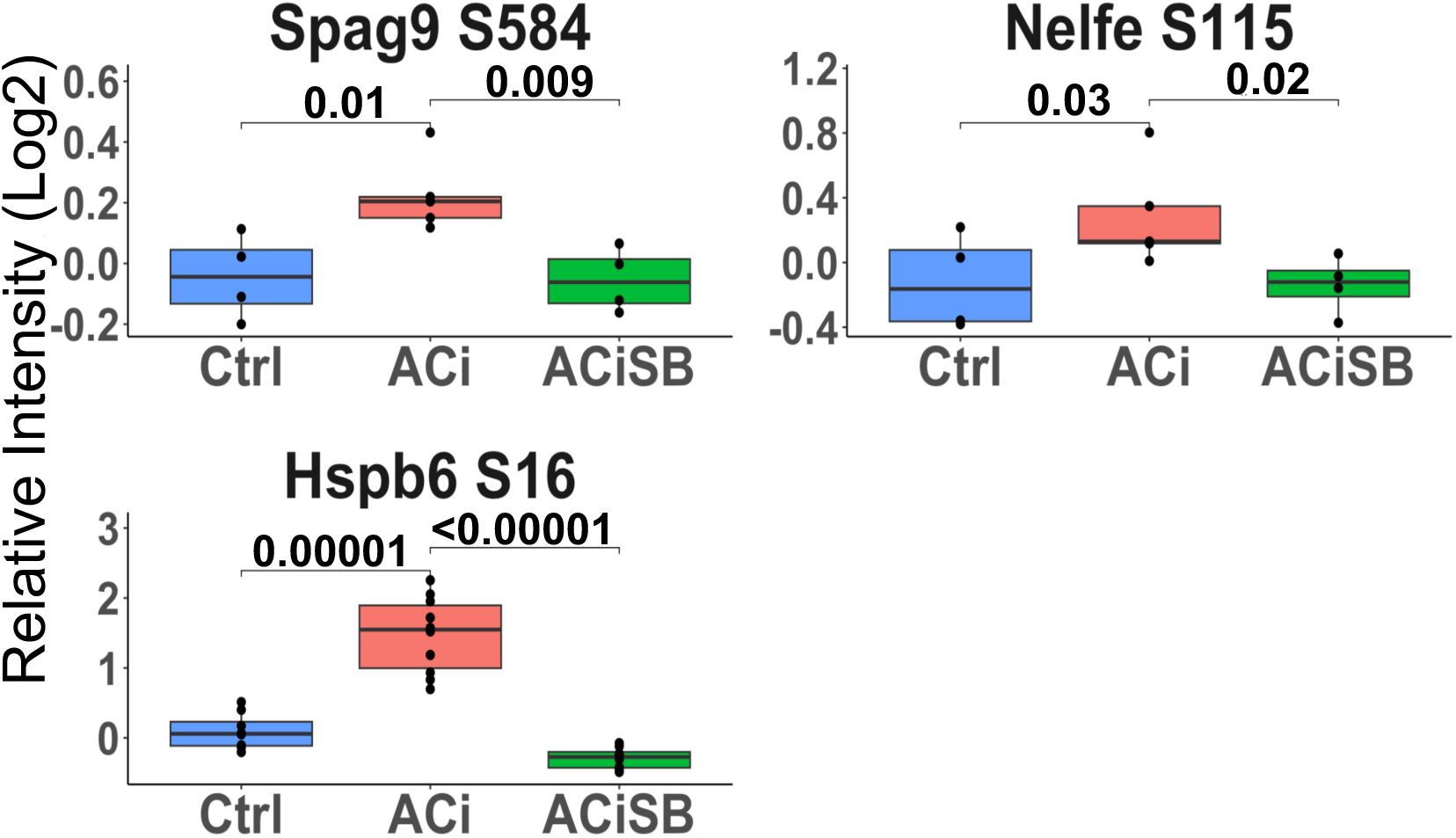
Phosphorylation of known p38 MAPK substrates and targets. Global significance (p < 0.05) established via F-test, with p-values derived from LIMMA contrast matrices for inter-group comparisons. P-values < 0.05 are indicated by faceted brackets.

**Figure S7.**
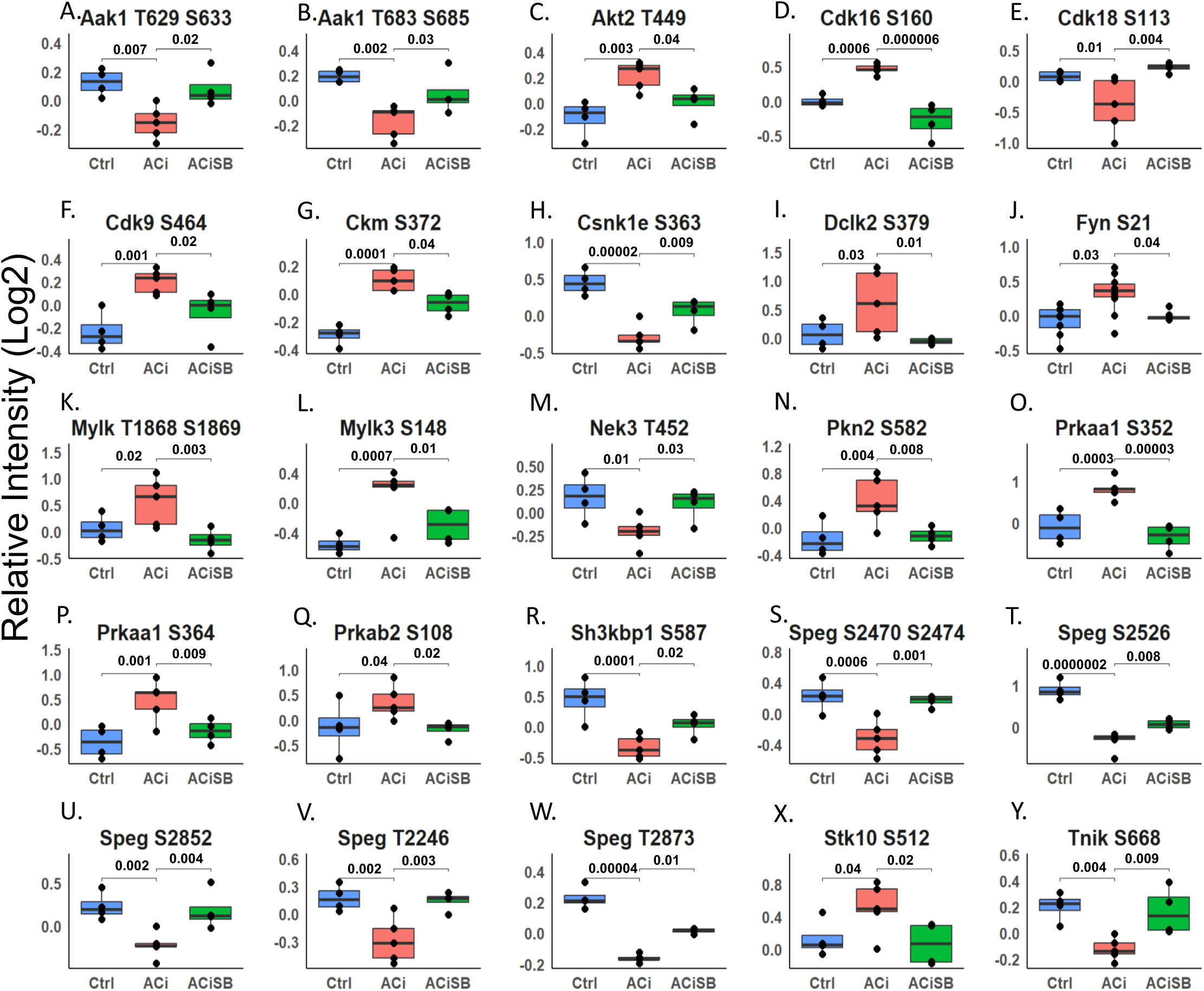
ACISB-Responsive Kinase Phosphorylation. Global significance (p < 0.05) established via F-test, with p-values derived from LIMMA contrast matrices for inter-group comparisons. Only a subset of results is shown in the figure. P-values < 0.05 are indicated by faceted brackets.

