## Supplemental File S1 - Methods for "Inhibition of Cardiac p38 Highlights the Role of the Phosphoproteome in Heart Failure Progression"

#### *Animal model*

Hartley guinea pigs (~300 g; HillTop Lab Animals) were housed in the animal facility at the Johns Hopkins University. This study conforms to the Guide for the Care and Use of Laboratory Animals published by the National Institutes of Health (NIH Publication No. 85-23, revised 1996) and was approved by the Johns Hopkins Animal Care and Use Committee. Male animals were anesthetized with 4% isoflurane in a closed box for 4 min, and then intubated. Animals were ventilated with oxygen and 2% isoflurane. Ascending aortic constriction (AC) was produced by tying a suture around the ascending aorta using an 18-gauge needle as a spacer, which was then removed. Sham operation was performed following the same procedure without the ligation.  Daily bolus of Isoproterenol (ISO) was administered via a programmable iPRECIO® pump (SMP-200, Data Science International, St. Paul MN) that was implanted in the peritoneal cavity. The pump was programmed for 1-hour daily delivery of Isoproterenol for a dose of 1 mg/kg/day. For the treatment of SB203580, an osmotic pump (Alzet Osmotic pump, model 2004, (Cupertino, CA) was implanted in the abdominal cavity. SB203580 was delivered continuously at 0.5 mg/kg/day DA. A vehicle filled osmotic pump was implanted for control in ACi group.  The following treatment group were studied:  1. Sham-operated, serving as controls 2. ACi (Aortic constriction +Isoproterenol treatment till end point)  3. ACi-SB (Aortic constriction +Isoproterenol treatment+ SB203580 treatment) All personnel involved in data collection and analysis were blinded to the treatment and non-treatment groups. Both groups received similar incisions and thus, could not be distinguished based on these interventions. Each animal was assigned a unique computer-generated numeric ID.

*Echocardiography*

Transthoracic echocardiography was performed on conscious non-anesthetized guinea pigs by using a Vevo 2100 high-resolution in vivo imaging system with 24 MHz transducer (VisualSonics, Toronto, ON, Canada) and analyzed with the Advanced cardiovascular package software (VisualSonics).

*Data and statistical analysis*

For heart weight, lung weight and FS% analysis between groups, one-way analysis of variance (ANOVA) followed by Tukey's post hoc analysis was used.

### *Proteomic Studies*

*Protein extraction and Immunoblotting*

Ventricles were harvested. Tissues were rinsed in cold PBS, rapidly heat-stabilized (Stabilizor$^{TM}$, Denator, Inc.), snap-frozen in liquid nitrogen, and stored in a -80 freezer. To extract protein, stabilized tissues were homogenized with RIPA buffer in the presence of 2% SDS, solubilized, and boiled in 1x LDS sample buffer for SDS-PAGE. The protein mixture was separated on a 4-12% NuPAGE gel (1 mm, Invitrogen). Samples were run at room temperature for 35 min at 200 V. Proteins were transferred to nitrocellulose membranes with iBlot (Invitrogen, Inc.), using program 3 for 7 min. Membranes were stained with Ponceau S solution (Sigma-Aldrich) to evaluate the transfer efficiency. Membranes were blocked for 1 h using Odyssey® blocking buffer (Li-Cor Biosciences) and incubated with primary antibody overnight at 4°C. Antibody binding was visualized with an infrared imaging system using IRDye secondary antibodies and quantification of band intensity was performed using the Odyssey Application Software 3.0.

| **Antibodies** | **Brand** | **Cat #** | **Species** | **Type** | **Dilution** |
|---|---|---|---|---|---|

| | | | | | |
|---|---|---|---|---|---|
| p38 MAPK Antibody | Cell Signaling Technology | 9212 | Rabbit | primary | 1:1,000 |
| Phospho-p38 MAPK (Thr180/Tyr182) (D3F9) XP® Rabbit mAb | Cell Signaling Technology | 4511 | Rabbit | primary | 1:1,000 |
| IRDye® 800CW Goat anti-Rabbit IgG Secondary Antibody | Li-Cor | 926-32211 | Goat | secondary | 1:10,000 |

Densitometric analysis of p38 and phosphorylated p38 (p-p38) signals in three distinct groups was conducted using Image J software, with data derived from digitized .tif image files. These signals were subsequently normalized to total protein levels detected via Ponceau staining. Statistical analysis was performed using one-way analysis of variance (ANOVA), followed by Tukey's post hoc test, to evaluate the comparison of p-p38/p38 ratios across the different groups.

*Sample Preparation, Proteolysis and TMT labeling*

The experiment compared 3 experimental groups: 1) Sham-operated controls, 2) ACi failing hearts (at 4 weeks) and 3) ACi animals treated concomitantly with SB203580 (0.5 mg/kg/day).

*Sample Preparation, Proteolysis and TMT labeling*

Hearts were harvested and washed with ice cold phosphate-buffered saline. The left ventricle free wall was isolated and immediately subjected to heat denaturation to abolish all enzyme activity using the Denator Stabilizor System (Denator) according to the manufacturer's instructions. Denatured samples were then stored at -80° C until the next step. The left ventricle of each guinea pig heart was homogenized with a handheld Polytron tissue disrupter in filtered and deionized, Tris-buffered 9M urea (5mL), pH 7.5 Samples were allowed to solubilize for 30 min at room temperature. Soluble homogenates were subjected to methanol/chloroform/water extraction and protein precipitation by the method of Wessel & Flugge(1). Samples were dried under nitrogen gas to remove residual chloroform before resolubilizing for 30 min in 9M urea. Aggregates were disrupted by brief bursts of sonication (< 30s total). Peptides were diluted 6-fold into 60 mM HEPES, 0.6 mM DTT, pH 7.5 such that the final reaction buffer contained 50 mM HEPES, 1.5 M urea, 0.5 mM DTT, pH 7.5. All samples were diluted further with reaction buffer to a common final protein concentration of 1mg/mL. Samples (1mg) were subjected to proteolytic digestion with proteomics grade Trypsin (1µg/100µg protein; Promega) at room temperature, overnight. The following morning, samples were supplemented with Trypsin (1µg/100µg protein; Promega). At that time, DTT was added to samples at a concentration of 5 mM and the digest was allowed to proceed for 1 hour prior to peptide alkylation by addition of iodoacetamide to a final concentration of 15 mM. Alkylation was allowed to proceed 1 hour at room temperature in the dark. Peptides were subsequently acidified by addition of trifluoroacetic acid to a final concentration 0.5% (v/v) and purified by solid phase extraction using SepPak tC18 cartridges (Waters) on a vacuum manifold. Purified peptides were eluted with 60% (v/v) acetonitrile in aqueous 0.1% (v/v) formic acid. Peptides were evaporated to dryness on an Eppendorf Vacufuge. Peptide samples were resolubilized in triethylammonium bicarbonate (TEAB) pH 8.5 and labeled with TMT reagents according to the manufacturer's instructions.

*Chromatography & Mass Spectrometry*

Following TMT labeling of individual samples, peptides were pooled and subjected to high-pH reversed-phase liquid chromatography (bRP-HPLC) (2), as detailed by Foster et al. (3). Briefly, samples were fractionated at 250ul/min on a Waters BEH C18 column with a gradient running from 10mM TEAB to 90% Acetonitrile, 10mM TEAB over 105 minutes. The fractions were concatenated into 24 fractions of which 20% of each was taken for the expression proteome determination. The remaining 80% was combined into 11 fractions and $TiO_2$ enrichment was performed to assess the phosphoproteome.

Following concatenation, samples were analyzed using a nanoAquity nanoLC system (Waters) interfaced with an Orbitrap Fusion Lumos Tribrid Mass Spectrometer

(ThermoFisher Scientific). Peptides were injected onto a 2-cm trap column at 5 µL/minute for 6 minutes before being eluted onto a 75 µm×15 cm in-house packed column (Michrom Magic C18AQ, 5 µm, 100Å) operating at 300 nL/min. Each sample was run on a 90-minute gradient. Data were acquired using a 3 second cycle between MS1 scans in FT-FT acquisition mode. The survey full-scan MS (400-1600 Th) was performed at a resolution 120,000 with an automatic gain control target ion intensity of $4x10^5$, while $MS^2$ scans were performed at a resolution of 50,000 with a target value of $1.25x10^5$. Mass tolerances for were +/- 10ppm for MS1. Maximum injection times were set to “Auto” for MS1 and 86 ms for $MS^2$. A 0.7 Dalton isolation window was used and HCD fragmentation was performed with a normalized collision energy of 36. Internal calibration was set to Easy-IC. Spectra whose charge was unassigned or +1 were not tabulated. Dynamic exclusion was set to a repeat count of 1 with a 15s exclusion time.

*Protein Identification*

The .Raw files for both the enriched and unenriched (12 + 24 respectively) were searched against a guinea pig database of predicted protein sequences (NCBI RefSeq, taxonomy: Cavia porcellus, date:01/20/2021, FASTA format, 37727 sequences), using Mascot Version: 2.8.0 (Matrix Science) interfaced through Proteome Discoverer 2.4.0.305 (Thermo). Peaks were filtered at a signal to noise ratio of 3, deisotoped, and searched with a parent ion mass tolerance of 5 ppm and an MS2 mass tolerance of 0.02 Da. Precursors that had co-isolation of 30% or greater were discarded from quantification. Trypsin was specified as the enzyme and 1 missed cleavage was allowed. N-terminal labeling with TMTpro reagent and carbamidomethyl were specified as a fixed modifications and dynamic modifications included deamidated NQ, oxidized M, phosphorylation on STY, and TMTpro on lysine. All searches were conducted with the reversed database search mode engaged. Percolator software was used for peptide FDR (q-value) calculations. Mascot output files (.dat) tabulated were in Proteome Discoverer. Only high confidence peptides only (q<0.01) were used for protein identification and quantification.

*Spectral Inclusion Criteria for Quantitation*

Analysis was confined to uniquely- and unambiguously assigned spectra (1% peptide FDR). Missingness of ion intensities for a single spectrum across reporter channels, was low (< 2% of unique spectra with intensities missing from one or more channels), indicative of efficient TMT labeling and fragmentation.

*Protein Quantification by TMT and Statistical Analysis*

Reporter ion intensities were integrated over 20 ppm using the most confident centroid method and corrected for purity in Proteome Discoverer 2.5 (Thermo). Spectral TMT signals were quantified using the median sweep algorithm originally described by Herbrich et al.(4) essentially as implemented recently by Foster et al.(3) with a minor modification. TMT reporter ion intensities were 1) logarithmically-transformed (base 2), 2) median-centered within each channel prior to 3) median-centering each individual spectrum across channels, 4) determining protein abundance by taking the median value of the logarithmically-transformed median-centered intensities for all spectra belonging to that protein in a given channel  (median summarization) For phosphopeptide analysis, all spectra (phosphorylated and unphosphorylated) median-centered, as above, and median summarized at the peptide level. Unphosphorylated peptides were filtered from the final table.

Following the median sweep, differential protein abundance between experimental groups was assessed using a Bayesian statistical framework, specifically, linear modeling of microarrays(5) (LIMMA) with multi-group comparison. Pairwise contrasts were also performed. The resulting moderated p-values were used to assess the positive false discovery rate (q-value) method of Storey (6, 7). The code used for the median sweep procedure and statistical analysis can be found at https://github.com/Frostman300/p38-upload. P-values derived from pairwise-comparisons to were utilized to determine specific differences between the groups, as demonstrated in the boxplots presented in the figures.

*Finding Homologous Human Peptides*

An Excel macro was written to find human peptides homologous to the detected guinea pig peptides using the command line version of Protein-Protein BLAST (2.15.0+). In brief, for each guinea pig peptide detected, blastp was used to compare the full length sequence of the corresponding guinea pig protein with a fasta format database of all Homo Sapiens refseq proteins downloaded from the NCBI protein database. Blastp was then used to align the guinea pig peptide with the corresponding highest scoring human protein target. The best homologous peptide match was then output to the spreadsheet. Macro code is available at https://github.com/Frostman300/p38-upload.

*Ingenuity Pathways Core Analysis Search Parameters*

For IPA core analysis of the proteome, uploaded data fields included the gene name and log2(ACi/ACiSB) value. Data were compared to the Ingenuity Knowledge Base reference set, which included genes only. The search space was limited to consideration of database relationships derived from primary tissues; data arising from cancer cell lines was excluded from analysis.Both direct(transcriptional ) and indirect(signaling) relationships were probed. Highlighted pathways were among those with Benjamin-Hochberg-corrected p-values of <0.05. Pathways are colored according to their z-score. The z-score integrates, not only gene over-representation, but the degree of concordance between supplied data (direction of change, magnitude, and p-value) and a database of curated relationships compiled from scientific literature and publicly-available datasets. IPA provides an inference about whether upstream gene-regulatory and signaling pathways may be activated or inhibited. Higher scores, whether positive or negative, reflect the strength of the inference(8).

*Network Analysis*

Functional protein association/interaction networks were constructed by loading the gene identifiers of up and downregulated proteins into stringApp 2.0.3 (9) embedded in

Cytoscape 3.10.2 (10). The default association/interaction threshold (STRING score > 0.4) was used to map relationships between proteins. Network modularity was assessed with the Markov clustering function in the clusterMaker2 app (v.2.3.4) (11) using the STRING score (>0.6) for edge weighting. The granularity parameter (inflation value) was set empirically. Modules were rearranged for clarity and named consistent with STRING's multipathway enrichment terms (e.g. Gene Ontology, Reactome, Kegg, among others). Singletons that were ontologically consistent with a module were grouped with it. The phosphorylation association network was further annotated using Omics Visualizer 1.3.1. (12) embedded in Cytoscape.
